# Multiple lines of evidence combine to understand tree species coexistence

**DOI:** 10.1101/2021.07.13.452199

**Authors:** Gordon G. McNickle, Morgan V. Ritzi, Kliffi M.S. Blackstone, John J. Couture, Taylor Nelson, Brady S. Hardiman, Madeline S. Montague, Douglass F. Jacobs

## Abstract

Understanding drivers of species coexistence is a central challenge in ecology (*1*, *2*). Coexistence cannot be observed directly, and while species co-occurrence in time and space is necessary for coexistence, it is not sufficient to prove coexistence (*3*, *4*). Species exclusion from a region is potentially observable, but can take decades to occur, and still might just occur stochastically (*5*, *6*). Thus, ecologists generally use theory to identify indirect observations that are indicative of mechanisms driving coexistence or exclusion (*7*–*9*). Various methods have been developed to indirectly infer coexistence, each of which requires different data, none of which are usually conclusive on their own, and are not typically combined (e.g. (*9*–*12*)). Here, we combine three different approaches which are normally used in isolation to build a case about a lack of coexistence of multiple hardwood species. Importantly, though each method has its flaws, we find that all three point to the same conclusion which strengthens the case for a lack of coexistence. First, in an experimental planting of three mature tree species we found no relationship between productivity and species diversity, which has been hypothesized to occur due to a lack of niche differences among species. Second, we used modern coexistence theory to calculate niche and fitness differences for each pair of species, which confirmed the high niche overlap among species and competitive dominance of one species. Third, we used the United States Department of Agriculture Forest Inventory and Analysis data to examine co-occurrence patterns of our species across thousands of natural forest stands and found that, these three species were distributed randomly throughout the USA, and do not significantly co-occur as would be expected if coexistence occurred naturally. Given that these independent methods agree; we take this as strong evidence about a lack of coexistence.

## MAIN TEXT

Ecological species coexistence has a well-developed body of theory (*7*, *8*, *13*), but is challenging to study empirically (*4*, *14*, *15*). Co-occurrence is necessary for coexistence but not sufficient to prove coexistence. For example, species may simply find themselves thrust together or apart for entirely stochastic neutral reasons (*3*, *4*, *7*). Ecologists therefore must rely on theory to identify experimental results that are indicative of coexistence as defined by theory (*9*, *10*, *16*). Generally, ecologists expect that differences among species in many dimensions (e.g. resource use, habitat use, seasonality) control coexistence such that there is some limit to how similar two species can be before one excludes the other. The totality of these dimensions, describing the differences among species, is referred to as the niche of a species (*1*). However, ecologists increasingly recognize the importance of fitness differences that might negate niche partitioning (*7*, *13*). Here, we combine multiple approaches that have been used over the years to infer species coexistence. On their own, each approach is inconclusive, but when all three approaches are combined they can complement the weakness of the others.

First, one of the oldest approaches to indirectly infer mechanisms of coexistence is to examine the relationship between community productivity and species richness (*5*, *17*). Generally, though not universally, researchers have found that more species rich plant communities are also more productive (*10*, *11*). One explanation for a positive diversity-productivity correlation would be if competition within a species was more intense than competition among species (*18*). Under this hypothesis, low diversity communities grow poorly due to strong negative intraspecific feedbacks (i.e. niche overlap), while species rich mixtures grow better because of diluted negative effects caused by niche partitioning among species. However, this is not the only explanation. Others have argued that positive diversity productivity relationships simply reflect sampling error with an increasing probability of selecting large productive species as diversity increases (*19*).

To examine the relationship between productivity and diversity we planted three species of hardwood trees in 2007: (*Prunus serotina* Ehrh., *Quercus rubra* L. and *Castanea dentata* (Marsh.) Borkh.) in all seven possible combinations of one, two or three species, crossed with two different planting densities (1m, or 2m between trees). The experiment has three replicate blocks, is located in west-central Indiana (40°26’41.9”N 87°01’46.4”W) within the historic range of these three species, and is now a closed canopy forest with trees approaching 10m in height and up to 20cm in diameter at breast height (Fig S1). In the summer and autumn of 2018 we took wood cores from the base of three individual trees of each species from each plot and used dendrochronology to estimate the abundance and productivity of species in these plots between 2007-2017. The tree cores give us the diameter of the base of the tree in each year, and we used this to calculate the cross-sectional area of the tree trunk in each year (i.e. basal area; BA). We used BA per metre squared as an estimate of population density in each year. The area of each annual growth ring is an estimate of productivity and is calculated as the BA of the tree trunk in year *t* minus the area in year *t* – 1, also known as the basal area increment (BAI). Wood is only one part of tree productivity, thus, to estimate leaf production in 2018 and 2019, we used cylindrical litter traps (28.6cm diameter). Finally, to estimate fine root production in 2019 we used an ingrowth core method. In autumn 2018, a cylindrical core (7.5cm diameter, 50cm deep) was removed from the forest floor, fine roots were sorted out and the soil replaced inside of a nylon mesh tube. After one year, the core was retrieved in 2019, roots washed, dried and weighed.

We found that the diversity and productivity of leaves, wood and roots were uncorrelated in these plots across all years (Fig 1; Table S1–3). This is somewhat rare, as positive relationships are overwhelmingly the most common in both manipulated biodiversity experiments and natural forest stands (*10*, *11*). Given contrasting interpretations of diversity productivity relationships, the lack of a positive relationship is either: (i) an indicator of a lack of niche differences among our species (*20*) or; (ii) simply evidence that our species grow to similar sizes at similar rates (*19*).

**Fig 1:**
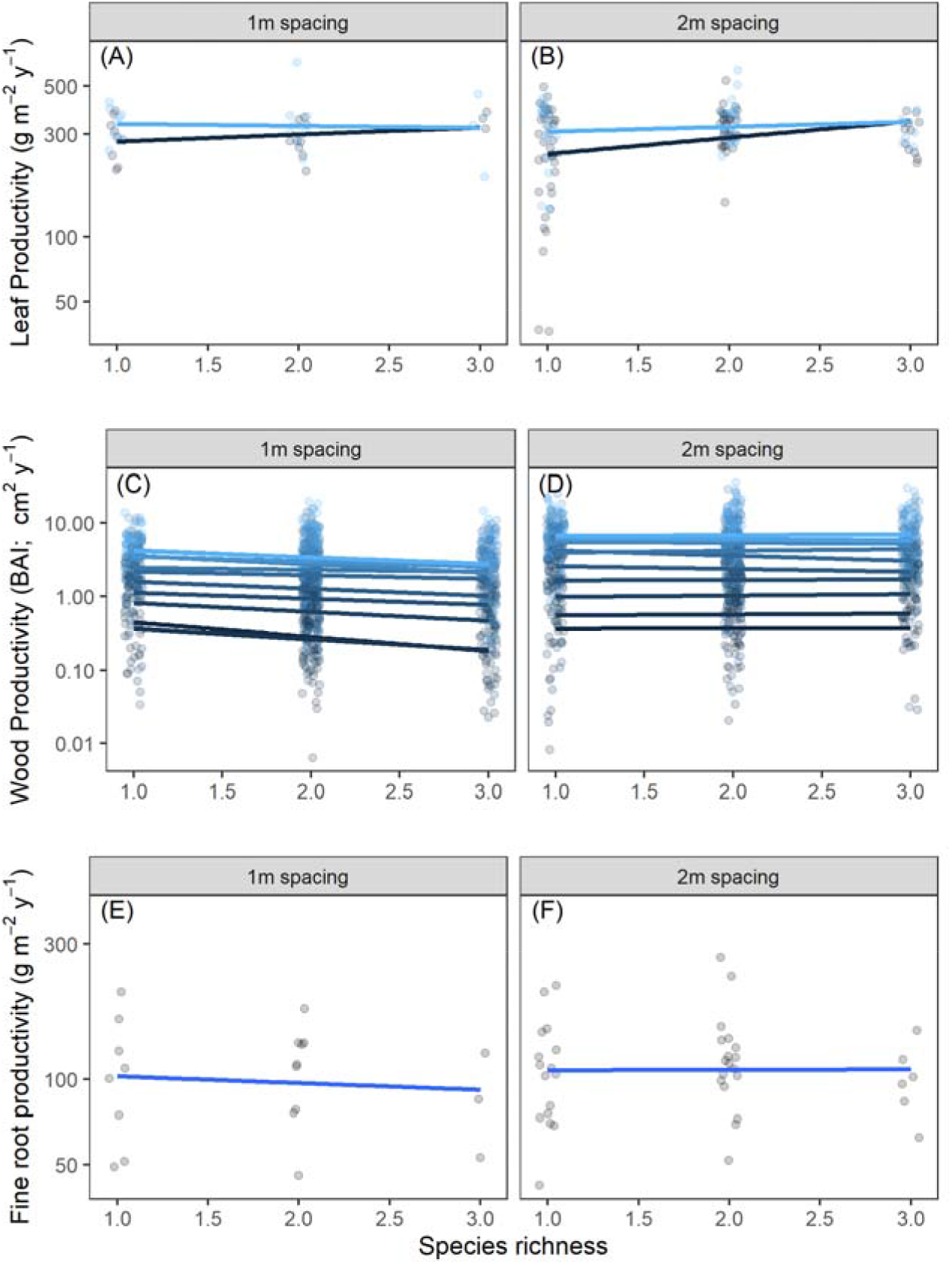
The relationship between species richness, and wood productivity estimated from leaf litter traps from 2018-2019 **(A,B)**; dendrochronology as basal area increments from 2007-2017 (BAI; **C,D**), and fine root productivity in 2019 (**E,F**). The y-axis is on a log_10_ scale in all panels. Trees were spaced either 1m or 2m apart (1 tree/m^2^, or 0.5 trees/m^2^ respectively). Points are drawn with a jitter around richness, and the y-axis is plotted on a log_10_ scale. Lines come from a linear regression (Tables S1–3). None of the estimates for the individual slopes of the diversity productivity relationship were different from zero.

To test these alternative explanations of the relationship between diversity and productivity, we used modern coexistence theory (MCT) (*7*, *13*). MCT focuses on the dynamics of populations, and partitions coexistence into two components: stabilizing mechanisms and fitness equalizing mechanisms (*7*, *13*). Stabilizing mechanisms of coexistence are mechanisms that cause one species to limit itself more than other species meaning that the negative effects of intraspecific interactions are stronger than any effects of interspecific interactions (*13*). Stabilizing mechanisms arise because of niche differences such as adaptation to different environments, or differences in biotic interactions (*13*). Second, fitness equalizing mechanisms are mechanisms that minimize average fitness differences among species (*13*). Large fitness differences among species can negate stabilizing effects and lead to competitive dominance. Therefore, we hypothesized that if the lack of a positive relationship between productivity and species richness to be an indicator of non-coexistence rather than sample error in species choice, then we hypothesized that MCT would predict exclusion.

To estimate niche and fitness differences we used Bayesian non-linear regression to fit a Lotka-Volterra (LV) model of the form:

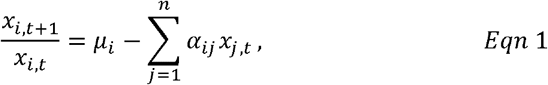

where *μ_i_* is the intrinsic rate of increase of species *i*; *α_ij_* is the competition coefficient for competition of species *j* on *i*; *n* is the number of competing species; *x_i_* is the population density of species *i*, and; *t* is the year in which *x_i_* was measured (Supplementary information). There are more than 10 definitions of niche and fitness differences that vary in their applicability to ecological interactions. We used the most inclusive definition for MCT that can account for facilitation and mutualism (*21*) where from Eqn 1, niche differences (ND) are given by:

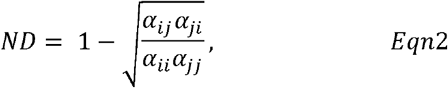

and fitness differences (FD) are given by:

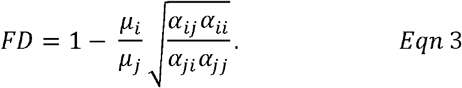

In the case of negative frequency dependence, regions of coexistence and exclusion can be demarcated by:

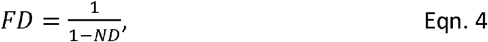

such that exclusion occurs above and to the right of this curve, and coexistence occurs below and to the left of the curve (*7*). In the case of positive frequency dependence (*22*), exclusion and priority effects are demarcated by:

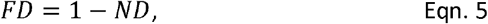

where exclusion occurs above and to the right of this curve, and priority effects below and to the left.

Population density trajectories suggest that *C. dentata* is competitively dominant and would eventually exclude the other two species (Fig 3A, B). Indeed the American chestnut was once dominant across forests in the northeastern USA before being almost completely extirpated by an invasive fungal pathogen (*Cryphonectria parasitica* (Murrill) M.E.Barr). Dominance of *C. dentata* is also visible from the MCT calculations which revealed that the niche differences among all combinations of the three species in our study were small, and fitness differences were such that competitive dominance led to exclusion rather than coexistence (Fig 2C; Table S4). These niche and fitness differences are consistent with the interpretation that the diversity productivity relationship was neutral because of niche overlap and not due to random sampling of species. However, generalizing would require information from natural stands across the native range of these species. Based on the diversity-productivity relationship (Fig 1) and MCT results (Fig 2), we hypothesized that these three species would not be found together more often than chance across their entire native range due to niche overlap, competitive dominance hierarchies or environmental filtering.

**Fig 2:**
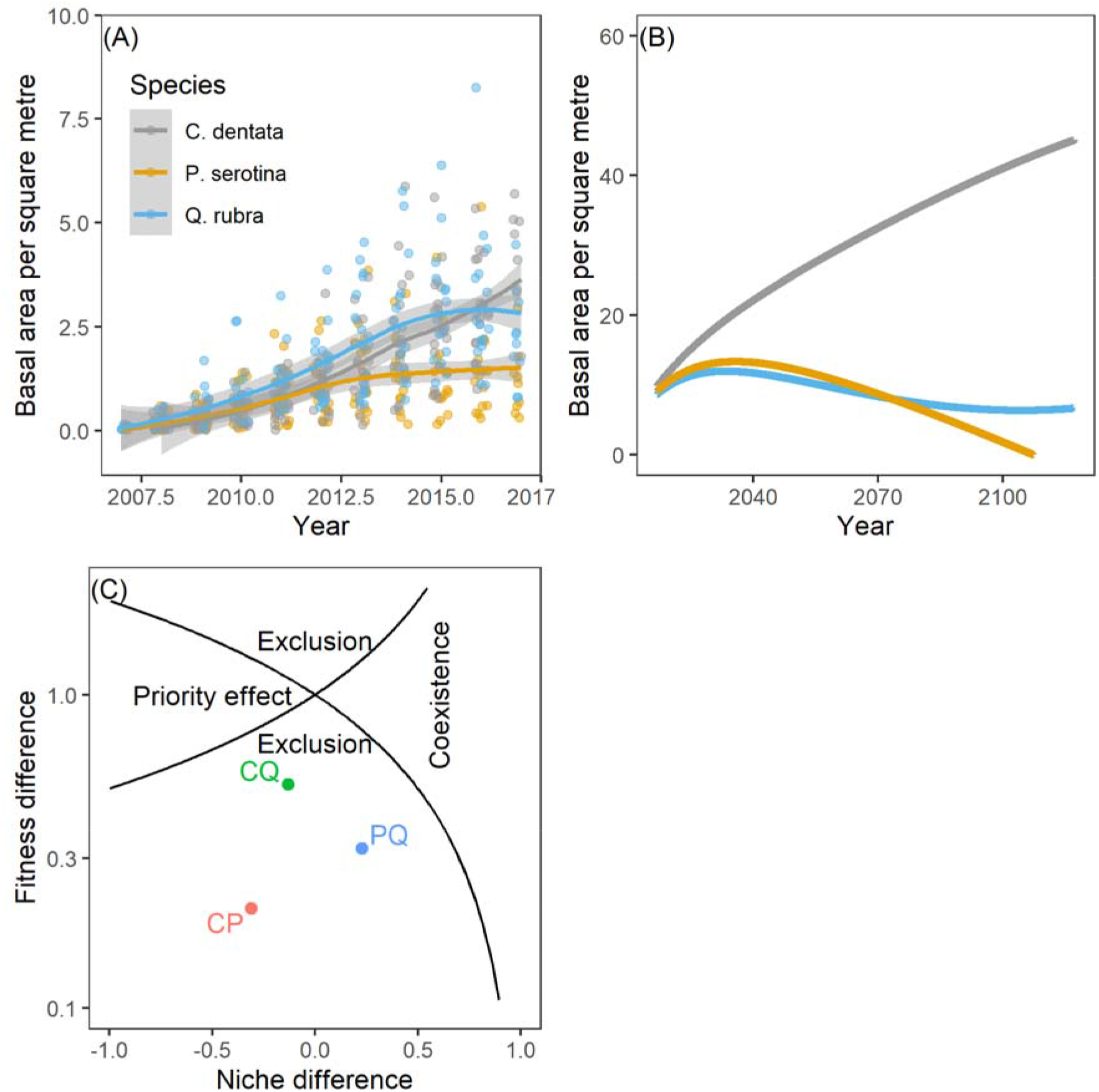
Population dynamics of the three species in the experimental plots. (A) Abundance through time estimated as basal area per m^2^. (B) Extrapolated population dynamics from numerically simulating 100 years of ODE behaviour from the estimated LV parameters. (C) The niche and fitness differences for each pair of species: *P. serotina* vs *Q. rubra* (PQ); *C. dentata* vs *P. serotina* (CP) and *C. dentata* vs *Q. rubra* (CQ). The black lines demarcate the regions of coexistence, priority effects and exclusion as labeled. The competitive dominance of *C. dentata* is evident in all three panels.

**Fig 3:**
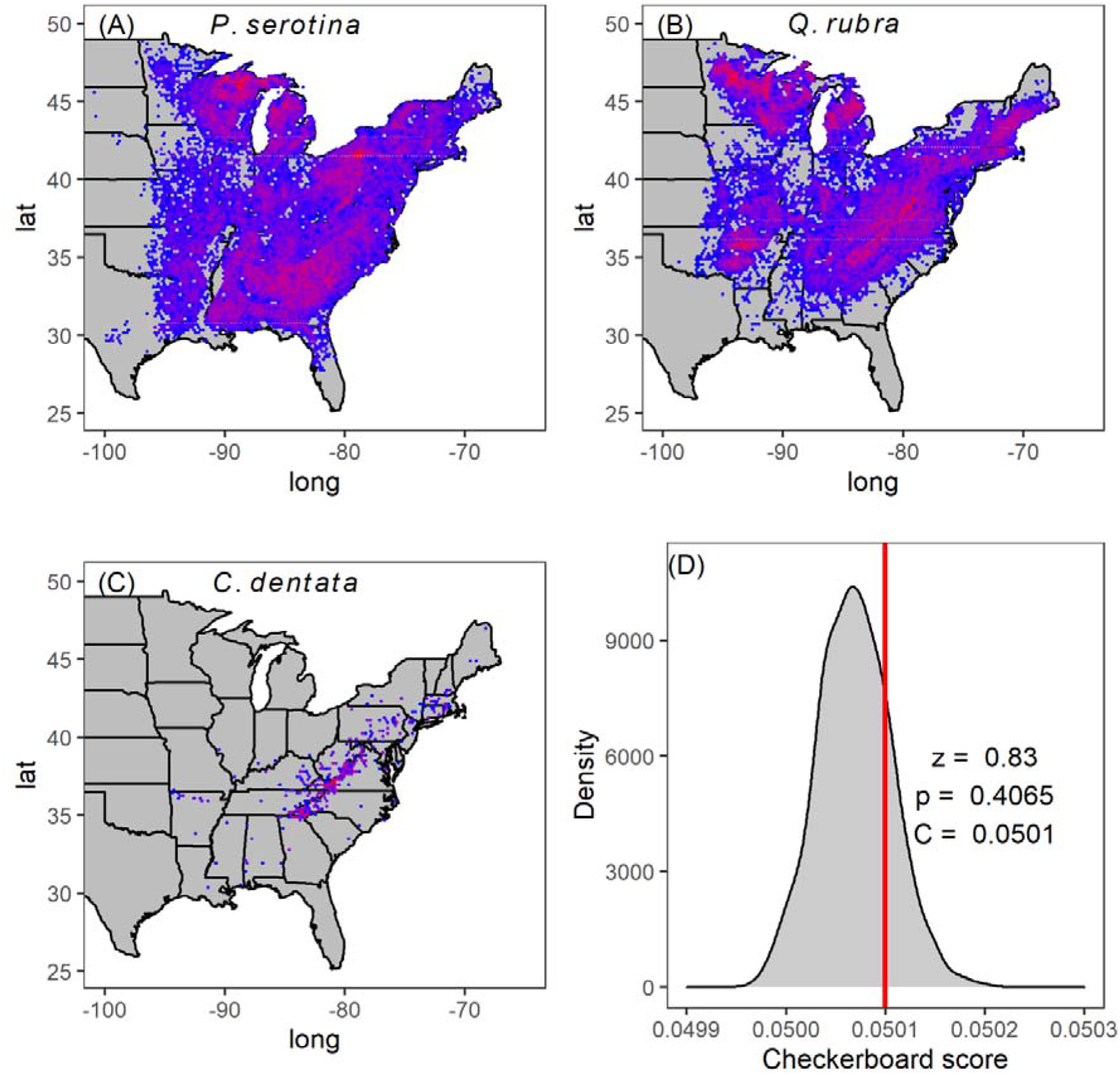
Distribution and population density (count) in natural forest stands of *P. serotina* **(A)** *Q. rubra* **(B)** and *C. dentata* **(C)** across the FIA plots in the USA. Red represents higher density, while blue represents low density. We compared observed C-score (red vertical line) with the null expectation of random distribution (grey histogram) across all FIA plots (D). The C score is high (relative to null) when species negatively co-occur, and low (relative to null) when they positively co-occur. The lack of difference compared to null (*p*=0.4065) suggests these three species are randomly distributed in nature with respect to each other on average.

To examine patterns of co-occurrence we used the publicly available USDA Forest Inventory and Analysis (FIA) data. The FIA data are an inventory of trees from thousands permanent 670 m^2^ plots with across the USA, including data for hundreds of thousands of trees allowing us to examine co-occurrence across the entire native range of our three focal species (Fig 3A-C). A wooded area of 670 m^2^ contains approximately 25 individual trees on average, and thus, is a reasonable scale to assess neighbourhood level competitive interactions. We used the checkerboard index (C-score) of co-occurrence (*3*, *23*, *24*) to quantify and statistically compare the observed co-occurrence pattern to a null expectation that the three species are randomly distributed throughout their range. C-scores that are significantly larger than the null indicate co-occurrence more often than chance which would be expected if these species frequently coexist in nature. We found that, at the scale of a 670 m^2^ FAI plot, the observed co-occurrence pattern was not significantly different from random (Fig 3D; *C_n_* = 0.0501, *z* = 0.83, *p* = 0.4065). This outcome means that, on average, these species do not positively co-occur throughout their range as might be expected if coexistence was common due either to niche partitioning or habitat filtering. A neutral distribution would be consistent with long-lived organisms at unknown stages of competitive exclusion. Checkerboard scores change based on the area sampled (*12*), and thus on its own the checkerboard score would be inconclusive. However, when combined with the weight of evidence from the lack of a diversity productivity relationship, and the niche and fitness differences estimated by MCT, we take this as further evidence that these species do not coexist at the scale of interaction among individual trees.

Here, by pulling on multiple threads of evidence, none of which was conclusive on its own, we have built a case that three species of hardwood trees cannot coexist. First, the lack of a positive relationship between productivity and species richness suggested a lack of niche differences among species (Fig 1), but on its own could be explained by something other than coexistence and thus may not be conclusive (*19*). Second, MCT independently confirmed the small niche differences and fitness differences indicative of competitive exclusion. However, these first analyses were from one stand of only 3500 trees. Third, we looked at co-occurrence patterns of these species across their entire native range. We found that the three study species randomly co-occur at the scale of a 670 m^2^ FIA plot other across their entire range (Fig 3). Random co-occurrence patterns would be expected in natural stands at unknown ages and stages of competitive exclusion. Each of these methods has been used separately to try and infer potential mechanisms of coexistence, but on their own each individual method is not always conclusive proof of coexistence. Predicting species coexistence has long been a goal of ecological theory, and the confluence of multiple threads of evidence all pointing in the same direction demonstrates that ecologists have made significant progress in uniting theory and observation surrounding species coexistence. Indeed, it was not a given that these three methods would agree to build a case about coexistence.

We suggest that going forward, combining multiple threads of evidence related to biomass production of monocultures and mixtures, demographic trajectories combined with modern coexistence theory to reveal fitness and niche differences as well as patterns of co-occurrence at large spatial scales can be a powerful set of tools that will better reveal patterns of ecological species coexistence. The first two approaches require manipulative experiments that vary both species diversity as well as population density, while the last requires observational data from natural systems. Data projects such as the USDA FIA data, or the US’ National Science Foundation National Earth Observatory Network (NSF NEON), or the Smithsonian’s ForestGEO project make such large scale patterns of co-occurrence increasingly easy to examine.

Understanding species coexistence, particularly for hardwood trees such as our study species, is not simply a problem for basic science. The world’s timber production increasingly relies on planted stands of trees. Planted forests grew from 5% to 7% of global forest cover between 2001 and 2010 and supply as much as 33% of global timber supply (*25*–*27*). Species diversity in forests is generally valued by society, considered to have conservation value for rare species, and can foster important wildlife resources for society (*28*). Thus, a greater understanding of how to successfully combine tree species into high diversity and high productivity timber stands also has significant applied science value too. Excitingly, understanding the mechanisms that drive tree species coexistence turns out to be one of those rare situations where basic and applied science, as well as economic, conservation and societal values are almost uniquely aligned.

## METHODS

### Experimental plot design

As described in the main text, we established an experimental planting of trees in west-central Indiana, USA (40°26’41.9”N, 87°01’46.4”W) in the spring of 2007. The main soil type is Rockfield silt loam, and the site is moderately well drained. Mean annual temperature is 10.4 °C and mean annual precipitation is 970mm (*29*). Three species of hardwood trees were planted: *Prunus serotina* Ehrh. (black cherry), *Quercus rubra* L. (northern red oak) and *Castanea dentata* (Marsh.) Borkh. (American chestnut) in all seven possible combinations of one, two or three species. These combinations were: (*1*) *P. serotina* monoculture, (*2*) *Q. rubra* monoculture, (*3*) *C. dentata* monoculture, (*4*) *P. serotina* and *Q. rubra* together, (*5*) *Q. rubra* and *C. dentata* together, (*6*) *P serotina* and *C. dentata* together, and (*7*) all three species together (Fig S1). In addition, we included three density treatments of 1m, 2m or 3m between trees which correspond to 10,000 stems per hectare, 2,500 stems per hectare, or 1,111 stems per hectare respectively. The experiment was arranged into a randomized split plot design where each replicate block contained three densities by seven combinations of species for a total of 21 plots per block. With three replicate blocks this was a total of 63 plots (Fig S1).

In 2007, trees were planted as bare-root seedlings. Seedlings were obtained from Cascade Forest Nursery in Cascade, Iowa, USA. The *Q. rubra* and *P. serotina* nursery seed source was from stands local to the nursery. Pure *C. dentata* seeds were collected from a forest stand near Galesville, Wisconsin, USA (*29*). Prior to planting, 2% glyphosate was applied to eliminate herbaceous vegetation. After planting, a 10-foot-high fence was erected around the perimeter of the entire experiment to prevent damage from deer. The fence was removed in 2012. In addition, a mixture of pendimethalin and glyphosate was applied in the rows between trees in 2008, 2009 and 2010 with backpack and side mounted band sprayer as needed to reduce competition from herbaceous weeds as the seedlings established. In 2011, the rows were mowed. Starting in 2012 herbicide and mowing was ceased because the trees were taller than herbaceous weeds.

Each individual plot always contained 56 trees arranged in a 7 tree by 8 grid (Fig S1). Thus, plot size depended on the density treatment. Plots planted with trees at a 1m spacing were 7 by 8 metres (54m^2^), plots planted with trees at 2m spacing were 14 by 16 metres (224 m^2^), and plots planted with trees at 3m spacing were 21 by 24 metres (504 m^2^). In total, the 63 plots take up approximately 2.4 hectares. In the mixed species plots, trees were planted sequentially across the row with seven trees in a torus wrap such that individuals of the same species were never planted next to each other (E.g. Fig S1B). The 26 perimeter trees around each plot were denoted as a buffer row to account for potential edge effects and were not measured, leaving 30 focal trees in the middle of each plot (Fig S1B). Thus, across the entire experiment there were 1,648 buffer trees and 1,800 focal trees for a total of 3,438 trees. Every one of the 1,800 focal trees were individually tagged and numbered.

### Experimental plot data collection

There was significant mortality between 2007 and 2017 in the southern most 3m spaced plots from a flooding event that occurred early in establishment (Fig S1C), and thus the 3m plots were dropped from most productivity analyses due to lack of replication. Wood production between 2007 and 2017 was estimated using dendrochronology on cores taken during the 2018 growing season. Of the 1800 total focal trees in the experiment, three of each species in each plot were randomly selected to be sampled for a total of 24 cores per species and 72 cores in total. Due to loss of replicates in the 3m spaced treatments, only the 1m and 2m spaced plots were sampled. Cores were air dried in the lab, mounted on wood blocks and sanded flat with sequentially finer sand-paper ending with 800 grit. Cores were imaged at a resolution of 1200 dots per inch. Ring widths on digitized images were measured in CooRecorder (v 9.00, Cybis Elektronik & Data AB, Saltsjöbaden, Sweden). Assuming the cross-sectional area of the trunk was a circle, we estimated basal area in each year. The area of a ring grown in year *t* was therefore estimated as the basal area at the end of year t, minus the basal area at the end of year *t* – 1 creating a basal area increment in each year. The data analysis included the fully factorial combination of year by species combination by density in a linear model (Table S2).

Starting in 2018, leaf litter was collected in buckets staked to the ground which were 28.6cm diameter, 36 cm deep, and had eight half inch holes drilled into the bottom for drainage. In 2018 and 2019, leaf litter collection began in September, and continued bi-weekly until December. Upon return to the lab, leaves were sorted by species, dried and weighed. Plot size differed by tree density, and so we applied sampling effort proportional to the plot size. This resulted in a grid of leaf litter traps spaced 6 metres apart across the site with one litter trap in the center of the 1m spaced plots, four in the center of the 2m spaced plots and nine in the center of the 3m plots. In 2018 we had a total of 258 sampling stations, as the four of the dead southernmost 3m plots were not sampled. Since sampling occurred every two weeks across three months, this generated 1,548 samples which had to be sorted to species. To ensure the most efficient data collection, using the 2018 data we determined that the number of litter traps could be reduced to 1 leaf litter sampling station in the 1m spaced plots, 2 in the 2m spaced plots and 3 in the 3m spaced plots with no change in estimated mean or variance at the plot level. Sampling stations to be removed were determined randomly. Thus, in 2019, only 114 leaf litter stations were sampled across three months in the autumn. Total leaf production in each year was analyzed by summing across the six sampling dates, and all species within each plot, divided by the area of the litter traps (0.064 m^2^). This calculation provided an estimate of leaf production per square metre per year. The data analysis included the fully factorial combination of year by species combination by density in a linear model (Table S1).

Fine root production in 2019 was estimated using root in-growth cores. In this method, a 7.5cm diameter and 50cm deep cylindrical soil core was extracted from the ground in the autumn of 2018. The living roots were sorted out of the soil and then the root-free soil was replaced inside a size 7 mesh cylinder. The in-growth cores were paired with litter trap stations, and so there was one in each 1m plot, two in each 2m spaced plot, and three in each 3m spaced plot for a total of 114 in-growth cores. After one year, in the autumn of 2019, we retrieved the in-growth cores. These were sorted into the top 10cm, the middle 20 cm, and the bottom 20cm, dried and weighed. Analyses summed all three depths within a plot and divided by the total surface area of the soil at the top of the core (0.0044 m^2^) to get an estimate of fine root production per year per square metre.

### Modern Coexistence theory

To estimate the parameters of the LV model, first, Eqn 1 was fit on monoculture data to estimate *μ_i_* and *a_ii_*. Second, using the monoculture estimates of *μ_i_* and *a_ii_* we fit Eqn 1 to each two species mixture to estimate the six different values of *a_ij_*. We used BA per square metre as an estimate of population density between 2007 and 2017. Models were fit using a non-linear Bayesian estimation procedure using the brms library (*30*, *31*) in the R statistical environment (*32*). Models were fit using four Markov chain Monte Carlo (MCMC) chains, 2500 burn-ins, 5,000 iterations per chain, resulting in 10,000 estimates for each posterior distribution. No thinning was used as thinning has been shown to have no detectable effects on MCMC simulation other than increased computing time (*33*). The prior was assumed to be a normal distribution centred on 1 for *μ_i_*, and 0.5 for *α_ij_* and a variance of 1 for both.

### United States Department of Agriculture Forest Inventory and Analysis Data

In 1999, the Farm Bill (PL 105-185) directed the Forest Service to begin annual surveys of the national inventory of tree resources. These USDA FIA data are publicly available. They represent surveys from 124,563 permanent plots across the entire USA allowing us to examine the co-occurrence of our three study species across their entire native range (*34*). An FIA plot is made up of four sub-plots with a radius of 7.3 m (24 feet) arranged in an equilateral triangle, with one subplot at the centre, and the other subplots at the corners of an equilateral triangle centred on the first subplot with a total plot area of 669.7 m^2^. The identity and size of all trees with a diameter of ≥5 inches (12.7 cm) at a height of 4.5 feet (1.37m) in each subplot are then recorded.

### Checkerboard score analysis

The checkerboard score is named for the appearance of a perfect negative co-occurrence matrix, which takes on the appearance of a checkerboard based on the diagonal arrangement of 0s and 1s. To calculate the C-score, first, the number of checkerboard units for each pair of species is given by:

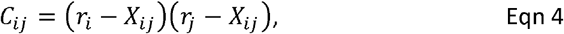

where *r_i_* is the number of times species *i* occurs without species *j, r_j_* is the number of times species *j* occurs without species *i* and *X_ij_* is the number of sites where species *i* and *j* co-occur (*3*). Typically, *C_ij_* is normalized by the number of possible species pairs (*P*) and the number of possible site pairs (*N*), which using basic combinatorics are given by

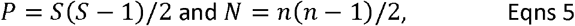

where S is the number of species, and *n* is the total number of replicate plots (*3*, *16*). Across all sites, the normalized C-score is given by:

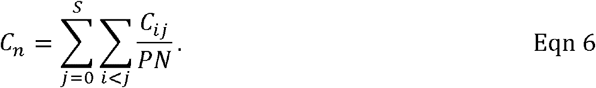

To generate the null hypothesis, we simply randomized the observed occurrence matrix from all FIA sites many times. The randomization procedure used a fixed-fixed randomization where row sums (i.e. species occurrences) and column sums (species occurrence within sites) are held constant (*35*). A fixed-fixed randomization is more conservative and less prone to Type-I errors. Null community matrices were created using the permatswap function in the vegan library in R (*36*, *37*). For each observed matrix, 1,000 null matrices were sampled with 30,000 swaps between each sample and with a burn-in of 30,000 swaps prior to the first sample. We used the quasiswap method for generating null matrices, which does not produce sequential null matrices but instead generates a matrix at each time step that is fully independent of previous matrices (*38*). The standardized effect size of the C_n_-score can be calculated based on the observed C_n_-score (C_n_) and the mean (μ) and standard deviation (σ) of the expected C_n_-scores produced from the 1000 null matrices by calculating a z statistic as (*24*, *39*):

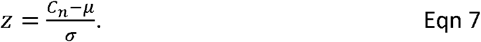

The *z* statistic is significant at the two-tailed p=0.05 level for −1.96<z>1.96.

## ACKNOWLEDGEMENTS

This work was funded by the American Chestnut Foundation, Purdue AgSeed, and a Hardwood Tree Improvement and Regeneration Centre grant to GGM, DFJ, JJC and BSH. Additionally, Hatch awards 1010722 to GGM and ND011490 to JJC, and McIntire Stennis project IND011535 to DFJ. The authors declare no conflicts of interest.

## AUTHOR CONTRIBUTIONS

MVR, KMSB and GGM performed dendrochronology analysis. MSM, DFJ and GGM performed root ingrowth core analyses. KMSB, TN, JJC and GGM performed leaf analyses. All authors contributed to tree tagging and survival data collection. GGM wrote the manuscript, performed MCT and C-score calculations and all authors contributed to revisions and discussion of approaches.

## DATA AVAILABILITY

The USDA FIA data are publically available from the USDA. Our site specific productivity and demography data are publically available on DRYAD at: https://doi.org/10.5061/dryad.280gb5mqc. All code required to reproduce analyses, figures and tables are publically available on Zenodo at: https://doi.org/10.5281/zenodo.5151988

## SUPPLEMENTARY INFORMATION

**TABLE S1:** Regression model for species diversity versus leaf production shown in Fig 1A,B. The model was lm(Leaf.production^~^ density * diversity * year) in R. The overall regression was not significant (F_7,160_ = 1.66, p=0.1217), and critically, none of the individual slopes for the diversity productivity relationship were significantly different from zero.

| Coefficients | Estimate | SE | t | p |
| --- | --- | --- | --- | --- |
| Intercept | 260.6 | 51.2 | 5.1 | <0.001 |
| Diversity | 21.3 | 27.7 | 0.77 | 0.443 |
| Density(2m) | -10.5 | 57.3 | -0.18 | 0.855 |
| Year(2019) | 76.2 | 72.4 | 1.0 | 0.295 |
| Diversity*Density(2m) | 6.0 | 30.9 | 0.19 | 0.847 |
| Diversity*Year(2019) | -19.8 | 39.1 | -0.51 | 0.614 |
| Density(2m)*Year(2019) | -13.0 | 84.9 | -0.15 | 0.88 |
| Diversity*Density(2m)*Year(2019) | 2.71 | 45.9 | 0.06 | 0.953 |

**TABLE S2:** Regression model for species diversity versus wood productivity shown in Fig 1C,D. The model was lm(BAI ~ density * diversity * year) in R. The regression was significant (F_43,1925_ = 24.1, p<0.0001), though the data contained a large amount of unexplained error (R^2^ = 0.33). None of the individual slopes for the diversity productivity relationship were significantly different from zero.

| Coefficients | Estimate | SE | t | p |
| --- | --- | --- | --- | --- |
| Intercept | 0.58 | 2.05 | 0.284 | 0.7761 |
| Density(2m) | 0.39 | 2.50 | 0.154 | 0.8773 |
| Year(2008) | 0.08 | 2.33 | 0.034 | 0.9728 |
| Year(2009) | 0.93 | 2.29 | 0.405 | 0.6859 |
| Year(2010) | 1.10 | 2.28 | 0.484 | 0.6283 |
| Year(2011) | 1.81 | 2.27 | 0.799 | 0.4246 |
| Year(2012) | 2.57 | 2.26 | 1.134 | 0.2567 |
| Year(2013) | 2.60 | 2.26 | 1.147 | 0.2516 |
| Year(2014) | 4.05 | 2.26 | 1.788 | 0.074 |
| Year(2015) | 4.29 | 2.26 | 1.892 | 0.0586 |
| Year(2016) | 5.24 | 2.26 | 2.314 | 0.0208 |
| Year(2017) | 4.79 | 2.26 | 2.114 | 0.0346 |
| Diversity | -0.08 | 0.94 | -0.087 | 0.9307 |
| Density(2m):Year(2008) | 0.28 | 2.96 | 0.094 | 0.925 |
| Density(2m):Year(2009) | 0.25 | 2.88 | 0.086 | 0.9312 |
| Density(2m):Year(2010) | 0.66 | 2.87 | 0.229 | 0.8193 |
| Density(2m):Year(2011) | 1.52 | 2.86 | 0.533 | 0.5943 |
| Density(2m):Year(2012) | 2.51 | 2.85 | 0.88 | 0.3787 |
| Density(2m):Year(2013) | 1.68 | 2.85 | 0.592 | 0.5541 |
| Density(2m):Year(2014) | 1.27 | 2.85 | 0.447 | 0.6546 |
| Density(2m):Year(2015) | 2.61 | 2.85 | 0.918 | 0.3585 |
| Density(2m):Year(2016) | 1.46 | 2.85 | 0.514 | 0.6076 |
| Density(2m):Year(2017) | 2.14 | 2.85 | 0.75 | 0.4531 |
| Diversity:Density(2m) | -0.08 | 1.16 | -0.066 | 0.9472 |
| Diversity:Year(2008) | -0.01 | 1.07 | -0.009 | 0.9931 |
| Diversity:Year(2009) | -0.17 | 1.05 | -0.162 | 0.8712 |
| Diversity:Year(2010) | -0.06 | 1.05 | -0.057 | 0.9542 |
| Diversity:Year(2011) | -0.25 | 1.04 | -0.239 | 0.8108 |
| Diversity:Year(2012) | -0.15 | 1.04 | -0.143 | 0.8866 |
| Diversity:Year(2013) | 0.03 | 1.04 | 0.026 | 0.9791 |
| Diversity:Year(2014) | -0.46 | 1.04 | -0.443 | 0.6576 |
| Diversity:Year(2015) | -0.40 | 1.04 | -0.38 | 0.7037 |
| Diversity:Year(2016) | -0.69 | 1.04 | -0.659 | 0.51 |
| Diversity:Year(2017) | -0.26 | 1.04 | -0.25 | 0.8025 |
| Diversity:Density(2m):Year(2008) | 0.02 | 1.38 | 0.011 | 0.9913 |
| Diversity:Density(2m):Year(2009) | 0.07 | 1.34 | 0.052 | 0.9587 |
| Diversity:Density(2m):Year(2010) | 0.12 | 1.34 | 0.09 | 0.9284 |
| Diversity:Density(2m):Year(2011) | -0.04 | 1.33 | -0.034 | 0.9727 |
| Diversity:Density(2m):Year(2012) | 0.39 | 1.33 | -0.295 | 0.7681 |
| Diversity:Density(2m):Year(2013) | 0.25 | 1.33 | 0.186 | 0.8523 |
| Diversity:Density(2m):Year(2014) | 0.81 | 1.33 | 0.614 | 0.5396 |
| Diversity:Density(2m):Year(2015) | 0.26 | 1.33 | 0.197 | 0.8438 |
| Diversity:Density(2m):Year(2016) | 0.80 | 1.33 | 0.606 | 0.5446 |
| Diversity:Density(2m):Year(2017) | 0.82 | 1.33 | 0.619 | 0.5358 |

**TABLE S3:**
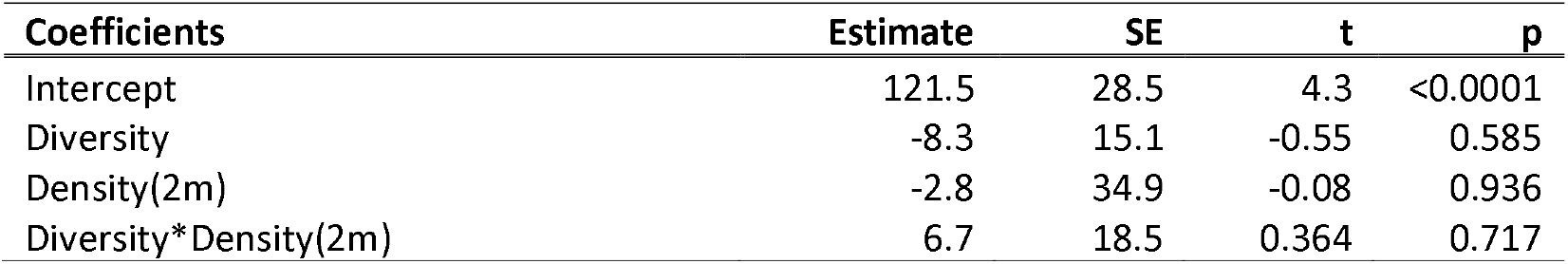
Regression model for species diversity versus root productivity data shown in Fig 1E,F. The model was lm(Root.production^~^ density * diversity) in R. The overall regression was not significant (F_3,56_ = 0.27, p=0.8476), and critically, none of the individual slopes for the diversity productivity relationship were significantly different from zero.

**TABLE S4:** Posterior results of Bayesian nonlinear estimation of LV model parameters. Species are abbreviated to the first letter of their genus as in all figures. Parameter and variable definitions can be found in the main text.

| Species $i$ | Species $j$ | Parameter | Estimate | Standard deviation | 95% credible interval |
| --- | --- | --- | --- | --- | --- |
| C | NA | $\mu_C$ | 1.3495 | 0.0658 | 1.2198, 1.4773 |
| Q | NA | $\mu_Q$ | 1.3780 | 0.0519 | 1.2764, 1.4783 |
| P | NA | $\mu_P$ | 1.2819 | 0.0671 | 1.1473, 1.4109 |
| C | C | $\alpha_{CC}$ | 0.0181 | 0.0052 | 0.0079, 0.0281 |
| C | Q | $\alpha_{CQ}$ | 0.0178 | 0.0046 | 0.0087, 0.0269 |
| C | P | $\alpha_{CP}$ | 0.0228 | 0.0085 | 0.0060, 0.0394 |
| Q | Q | $\alpha_{QQ}$ | 0.0331 | 0.0052 | 0.0228, 0.0433 |
| Q | C | $\alpha_{QC}$ | 0.0433 | 0.0049 | 0.0337, 0.0528 |
| Q | P | $\alpha_{QP}$ | 0.0118 | 0.0059 | 0.00002, 0.0231 |
| P | P | $\alpha_{PP}$ | 0.0209 | 0.0054 | 0.0100, 0.0315 |
| P | C | $\alpha_{PC}$ | 0.0284 | 0.0029 | 0.0228, 0.0342 |
| P | Q | $\alpha_{PQ}$ | 0.0352 | 0.0039 | 0.0274, 0.0429 |

**FIG S1:**
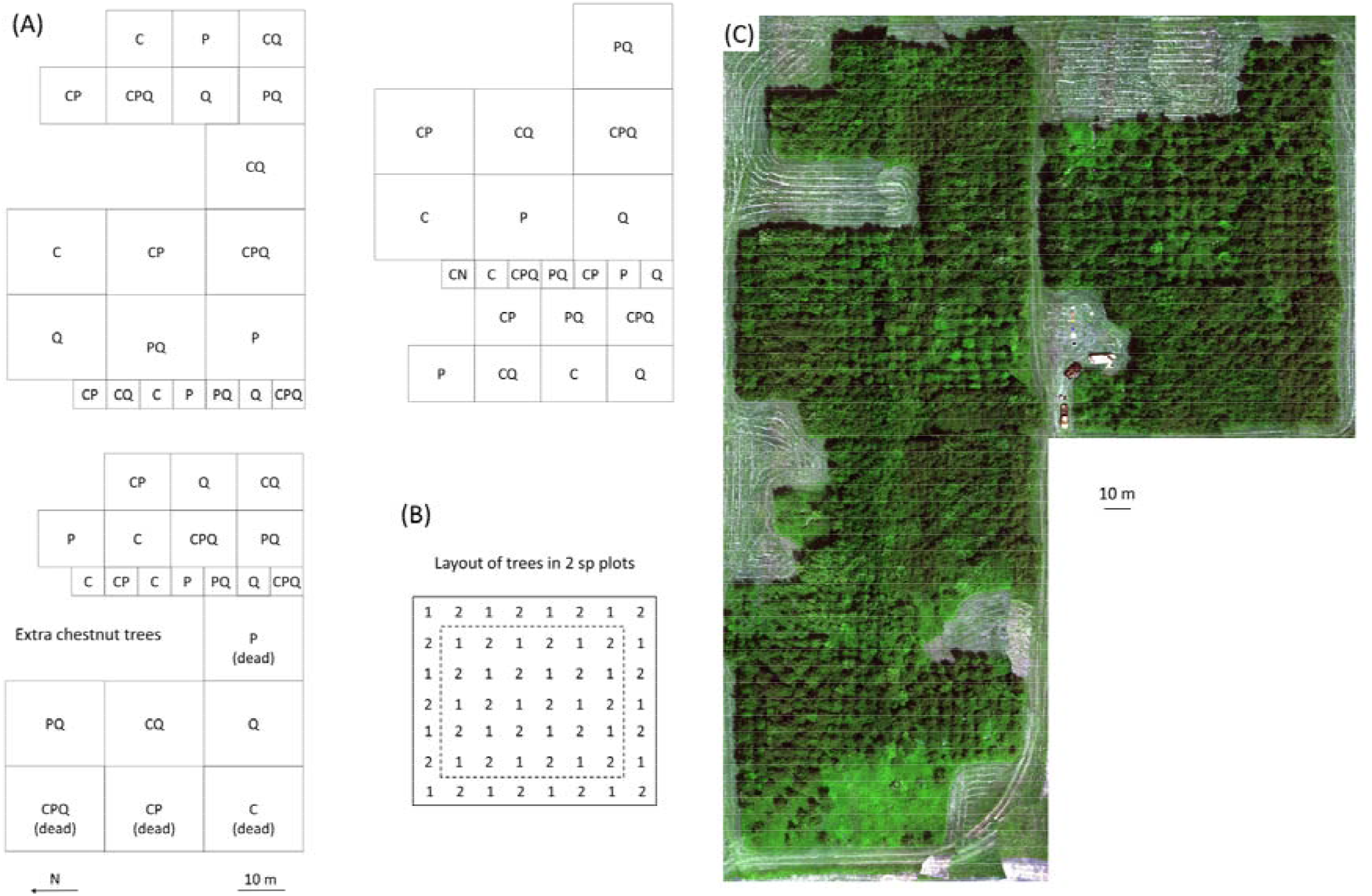
Plot layout. **(A)** The experiment has three replicate blocks that include all possible combinations of *C. dentata* (C, American chestnut), *P. serotina* (P, black cherry) or *Q. rubra* (Q, northern red oak), at three different planting densities (1m, 2m or 3m between trees; 1 tree/m^2^, 0.5 trees/m^2^ or 0.33 trees/m^2^ respectively) in a full factorial split plot design. Unfortunately, many trees in the southern most 3m spacing plots died due to an unusual flooding event. These plots are not included in productivity data due to inadequate sample size for analysis, however these plots are included in demography data. Surplus chestnut trees were planted in an open space at the southern end of the experiment because of their conservation value. **(B)** Each plot contains 56 trees that alternate between species (indicated by 1 and 2). There is a buffer row of trees that are not measured around each plot (dashed line) leaving 30 focal trees in the center of each plot, and 1260 individual focal trees in 1m and 2m spaced plots. (C) An aerial image of our plots showing the basic layout. The dead 3m spaced plots, and the extra chestnut trees that are not part of the experiment, are especially evident in this image. Field vehicles are also visible in the center of the image. Full methodological details about the planting and experimental design can be found in: (*29*).

